# Indirect Reciprocity Undermines Indirect Reciprocity Destabilizing Large-Scale Cooperation

**DOI:** 10.1101/2023.11.07.566116

**Authors:** Eric Schnell, Michael Muthukrishna

**Affiliations:** London School of Economics and Political Science (LSE)

## Abstract

Previous models suggest that indirect reciprocity (reputation) can stabilize large-scale human cooperation^1^. The logic behind these models^2–7^ and experiments^6,8^ is that a strategy in which individuals conditionally aid others based on their reputation for engaging in costly cooperative behavior serves as a punishment that incentivizes large-scale cooperation without the second-order free-rider problem. However, these models and experiments fail to account for individuals belonging to multiple groups with reputations that can be in conflict. Here we extend these models such that individuals belong to a smaller, “local” group embedded within a larger, “global” group. This introduces competing strategies for conditionally aiding others based on their cooperative behavior in the local or global group. Our analyses reveal that the reputation for cooperation in the smaller local group can undermine cooperation in the larger global group, even when the theoretical maximum payoffs are higher in the larger global group. This model reveals that indirect reciprocity alone is insufficient for stabilizing large-scale human cooperation because cooperation at one scale can be considered defection at another. These results deepen the puzzle of large-scale human cooperation.

---

Human cooperation takes many forms, occurring at different scales in different societies, multiple scales within the same society, and across different domains^9^. In some societies people primarily cooperate with extended families^10,11^ and in others cooperation occurs across large nation states and diverse ethnic populations^12,13^. The coexistence of scales can create problems such as corruption and nepotism, which can be interpreted as small-scale cooperation with family and friends undermining large-scale cooperation with society as a whole^14,15^. Indirect reciprocity has been proposed as a mechanism for aligning incentives to cooperate across multiple scales.

In a seminal model, Panchanathan and Boyd^1^ model a two-step cooperative game, where players play a Public Goods Game (PGG) followed by a Mutual Aid Game (MAG). In the PGG, players can apportion some of their endowment to a public good which is multiplied and then divided evenly among all players regardless of contribution. The multiplier (*M*) is less than the number of players (*N; M < N*), such that the Nash equilibrium is to contribute nothing to the public good. Players are then paired with group members at random and are given the choice to provide them with aid, which costs the provider and only benefits the receiver. The model reveals that an evolutionary stable strategy is for players to conditionally aid those who cooperate in the PGG and not those who defect. In this way, the MAG serves as a way to reward cooperators and punish defectors. Thus, indirect reciprocity – reputation for cooperation in the PGG - can maintain large-scale cooperation without the second-order free-rider problem. Later experiments support the insights from this model^8^.

Panchanathan and Boyd’s^1^ model and other similar models of indirect reciprocity^3,4,6,16,17^ fail to account for the existence of multiple possible cooperative reputations because people belong to multiple groups – multiple public goods games. For example, one could donate money to a local conservation group maintaining local parks or to a national or even international conservation group. In either case, this individual is donating to a cause that benefits others, but the scale of the cause and the circle of effected individuals differs. As an outside observer, how do we weigh these different actions and which do we choose to prioritize for improving a person’s reputation^18,19^? If cooperation at the local level is continuously prioritized over cooperation at the global level, this risks eroding large scale cooperation. This can be thought of as corruption, where small subsets of a larger group cooperate together at the detriment of the larger group. Existing models of indirect reciprocity fail to consider the implications of how different scales of cooperation interact on whether these scales can be aligned.

Here we extend Panchanathan and Boyd’s^1^ model such that players belong to multiple groups. In our model, players belong to one of *a* groups, which we call *local groups*. These groups and all players also belong to a larger cooperative group, which we call the *global group*. Local groups are all the same size and independent of one another. Players play two simultaneous Public Goods Games (PGG) where they can contribute to the local PGG, the global PGG or defect. Players then play a Mutual Aid Game (MAG) with a randomly chosen member of their local group. We analyze evolutionarily stable strategies using an adaptive dynamics approach, testing invasibility of 15 possible strategies against each other for a single rare mutant of each strategy and a group of individuals with each strategy. These 15 strategies are composed of 3 possible PGG strategies and 5 possible MAG strategies.

For the PGG players can either always cooperate in the global PGG (*G*), always cooperate in the local PGG (*L*), or defect, cooperating in neither (*D*). For the MAG players can either always provide aid (*c*), provide aid only to global PGG cooperators (*g*), provide aid only to local PGG cooperators (*l*), provide aid to others who aid in the MAG (*m*), provide aid to those who cooperated in the PGG and provided aid in the MAG (*pm*), or never provide aid to anyone (*d*). An example of how some of these strategies interact is shown in Figure 1. For completeness, in the appendix we also analyze PGG strategies which are dependent on having received aid in the MAG, but these analyses show no qualitatively different results.

**Figure 1:**
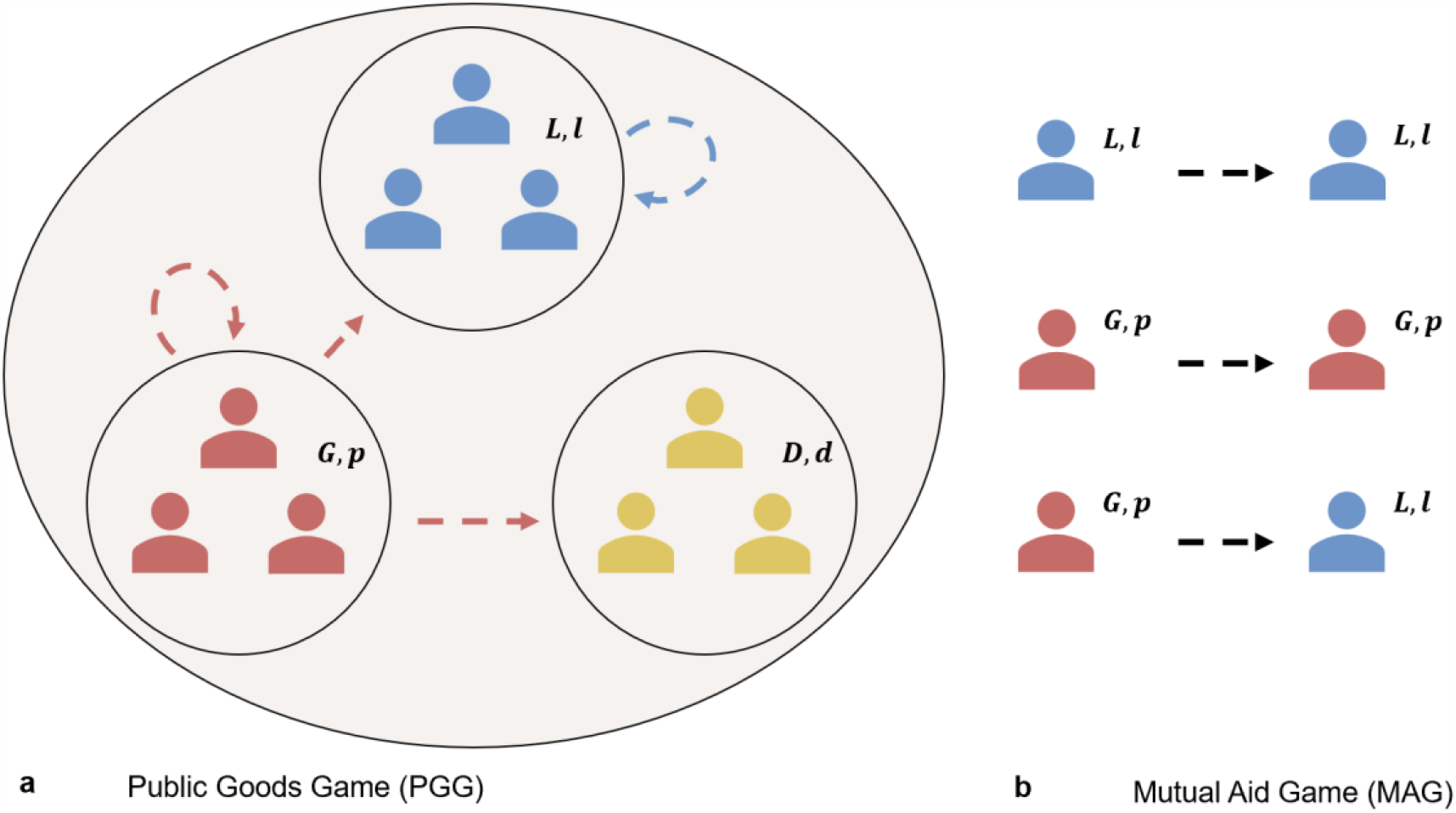
(**a**) Shows a perspective payout from different PGG strategies. In this example, players using strategies (*G, p*), (*L, l*) and (*D, d*) are each found in their own group with arrows showing where PGG returns go are distributed. (**b**) Shows which pairings from (**a**) will provide aid in the MAG. In this example, players using (*G, p*) help those who provided to either PGG and so aid players using (*L, l*), however (*L, l*) only provides aid to local cooperators and so won’t aid (*G, p*) in return. Crucially, this presents a second order free-rider problem, someone who cooperates in the first instance but won’t provide aid to cooperators in the second instance, showing the need for MAG strategies such as pm.

Consistent with previous models of large-scale cooperation sustained by indirect reciprocity, when there is effectively only one PGG and MAG (i.e. strategies (*G, pm*) and (*L, pm*)), indirect reciprocity is sufficient to sustain cooperation. That is, defectors (i.e. (*G, d*), (*L, d*), and (*D, d*)) cannot invade these strategies. However, when there are multiple scales of cooperation, then the smaller scale is more stable even when the multiplier and potential payoffs are higher in the global PGG, because fewer people need to cooperate in the local PGG. Conflicting reputations lead to smaller scale, local cooperation undermining larger scale, global cooperation.

We first analyze whether a single rare mutant of each strategy can invade a resident population of each other strategy. Consistent with Panchanathan and Boyd^1^, when there’s only one rare invader, defectors – PGG (*D*) and either defect (*d*) or reciprocate (*m*) in the MAG – can invade all other strategies, except for cooperative strategies utilizing indirect reciprocity, such as (G, g) or (*G, pm*). Such cooperative strategies utilizing indirect reciprocity are resistant to direct invasion by defectors but can’t invade these defectors in return. Furthermore, a reciprocal strategy that only cares about one of the PGG and MAG, such as (*G, g*) or (*G, m*), can be invaded by defectors in the long run through intermediate strategies which continue to cooperate in the PGG. This invasion pathway is shown in Figure 2. To prevent invasion of defector strategies, a resident population of PGG cooperators must have an MAG strategy that discriminates both on previous PGG and MAG behaviour when deciding who to aid, such as (*G, pm*) or (*L, pm*). That is, strategies in which players fall out of good standing if they’ve defected from either the PGG or MAG. These cooperative strategies while resistant to invasion from rare defectors also can’t invade a resident population of defectors. Thus, indirect reciprocity can ensure cooperators resist invasion from defectors, but a rare mutation of cooperation is not a solution how cooperation could evolve in the first place^20^.

**Figure 2:**
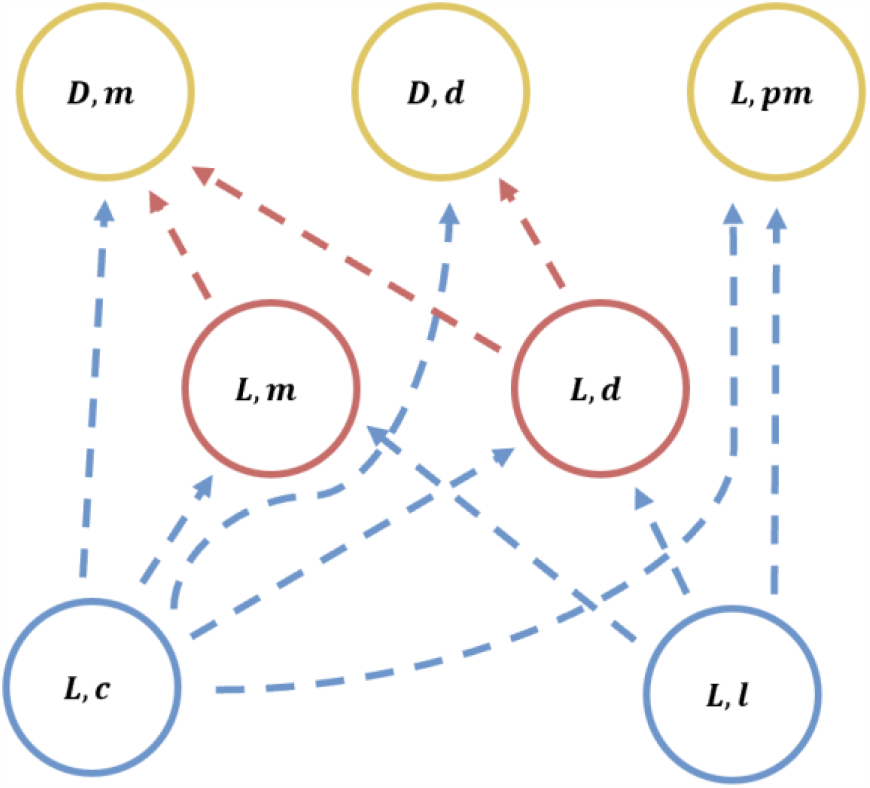
Example of invasion pathway for a single invader for parameters (*c*_*p*_ = 1; *c*_*m*_ = 1; *b*_*m*_ = 3; *b*_*g*_ = 3; *b*_*l*_ = 2; *n*_*l*_ = 5; a = 5; *e* = 0.05; *β* = 0). For clarity we’ve highlighted only one scale of cooperation (local), but the global scale behaves the same.

Cooperative benefits only emerge when there is more than one cooperator. This means, for cooperative strategies to invade others, mutants must invade in groups. As such, we next consider some percentage *β* of players in one local group with an invading strategy, with the rest of the resident local group (1 − *β*) as well as all other local groups using a resident strategy. We start by analysing a specific case of this approach where *β* = 1, or the entirety of one local group uses the invading strategy. Our analyses reveal that defecting strategies can only invade global cooperator strategies, not local cooperator strategies, and can only do so when:

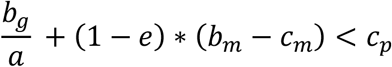

In other words, defectors can invade if the cost of cooperating in the PGG outweigh both the PGG benefits and net benefits from aiding and being aided in the MAG. This invasion pathway is shown in Figure 3. The reason defectors can invade at all is that they continue to free ride on the benefits provided by global cooperators in other local groups. For this same reason, defectors can’t invade local cooperators, because they don’t receive any benefits from the other local groups.

**Figure 3:**
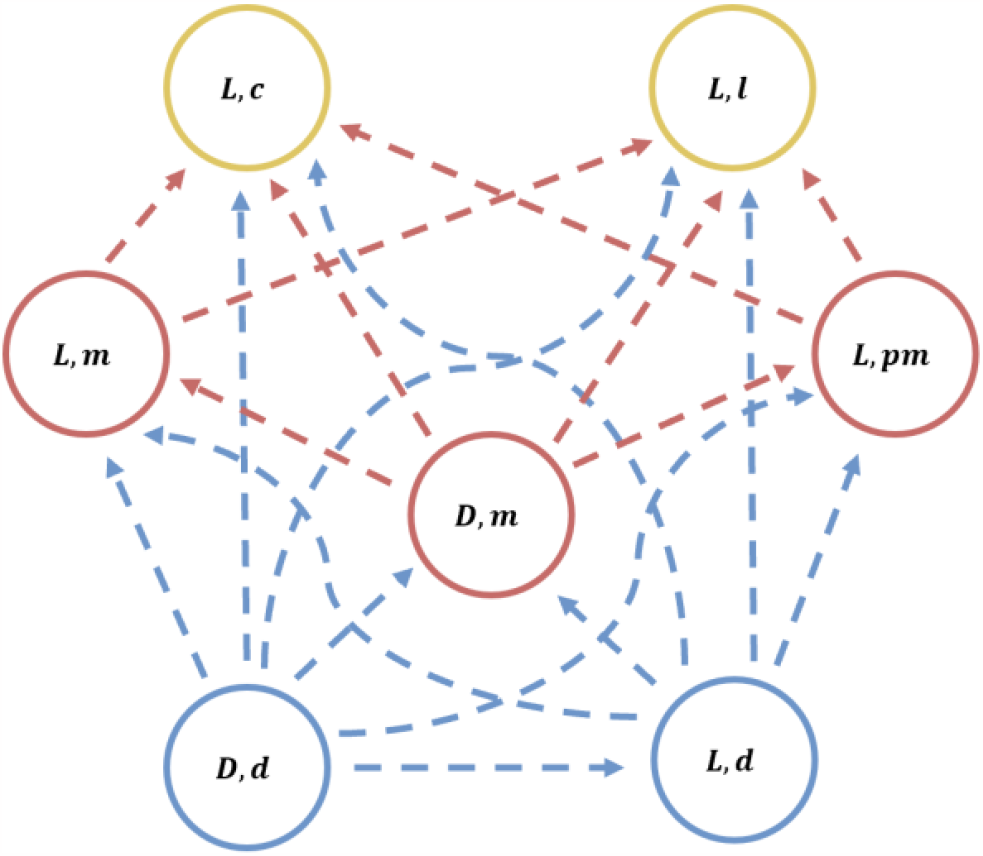
Example of invasion pathway for a group of invaders for parameters (*c*_*p*_ = 1; *c*_*m*_ = 1; *b*_*m*_ = 3; *b*_*g*_ = 3; *b*_*l*_ = 2; *n*_*l*_ = 5; a = 5; *e* = 0.05; *β* = 1). For clarity we’ve highlighted only one scale of cooperation (local), but the global scale behaves the same.

Local cooperators will invade global cooperators as long as:

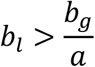

That is, for global cooperation to outcompete local cooperation requires benefits to higher than the benefits of all local groups combined (i.e. *a* · *b*_*l*_). Unless global cooperators can support all other local groups cooperating at a different scale, then local cooperation will undermine global cooperation even when the benefits of cooperating at a global scale are higher. Moreover, the more splintered the society is – the more local groups there are – the harder it is for global cooperation to outcompete local cooperation. This makes local cooperation much more stable and likely to emerge than global cooperation under a range of realistic parameters.

When *β*<1, defectors can begin to invade and dominate global cooperators when:

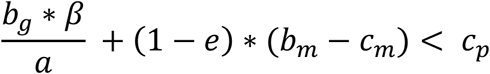

And they can invade local cooperators when:

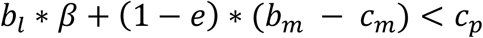

This is similar to the instance where invaders occupy a whole group, but the benefits of the PGG are lessened because there are fewer players contributing to it. As *β* reduces, the model resembles the single rare mutant analysis and cooperation is at a larger disadvantage. As Figure 4 reveals, as the number of cooperative invaders increases, the required returns from cooperation can be lower for cooperators to invade.

**Figure 4:**
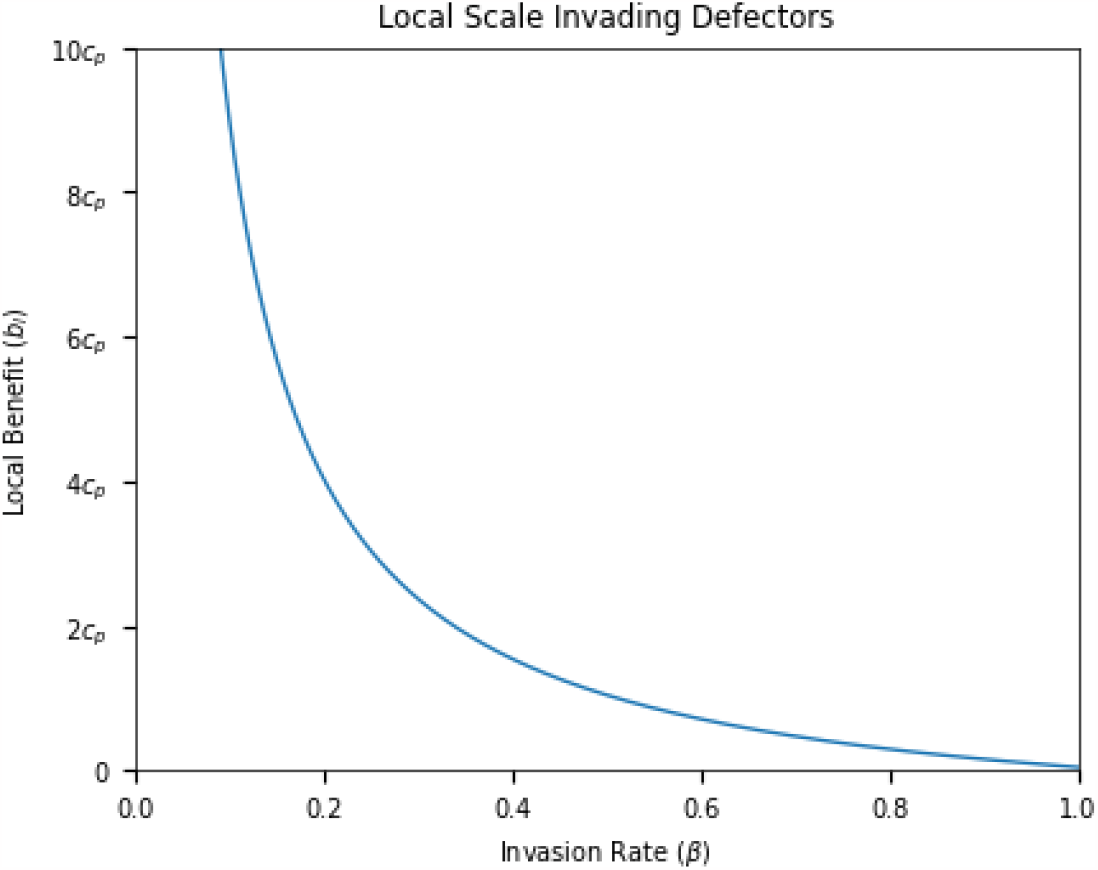
Relationship between the benefit of local cooperation (*b*_*l*_) compared to the cost of cooperating (*c*_*p*_) required for local cooperators to invade a defector resident, given a relatedness of invaders (*β*). As the number of mutants using a shared strategy increases, the required benefits of cooperation needed for invasion decreases. Parameters shown: *c*_*p*_ = 1, *c*_*m*_ = 1, *b*_*m*_ = 2, *e* = 0.05.

In all such cases, local cooperation still outcompetes global cooperation even with lower local benefits. Global cooperators can only invade local cooperators when:

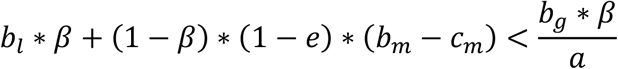

Thus, under a wide range of conditions and arguably all realistic conditions, indirect reciprocity when there is more than one possible reputation leads to lower scales of cooperation undermining higher scales. Global cooperation can only dominate when the global PGG benefit is so high that there is effectively no longer a dilemma between contributing to the local and global group. Thus, indirect reciprocity alone is not sufficient to sustain large-scale human cooperation.

Individuals rely on the reputation of others when making cooperative decisions^21–27^. But often these reputations can be in conflict^3,28,29^. The same person may have a positive reputation with some groups and a negative reputation with others in the same society; the same person may have a positive reputation in some domains and a negative reputation in others. This makes it difficult to determine with whom you should cooperate. Given that societies are made up of overlapping and embedded groups of differing sizes and scales of cooperation, it is necessary to reconcile reputational differences across different scales of cooperation.

This model shows how different scales of cooperation interact revealing that small-scale cooperation in a local group is more likely to be sustained than large-scale cooperation, even when cooperation is more beneficial at the larger scale. These results add further insight to an emerging literature on intergroup interactions. If group members interact more frequently across local group boundaries and move between groups, the effective population becomes closer to the global population incentivizing higher scale cooperation^6,30^. By corollary, if individuals are more likely to interact within a local group (e.g. local region or ethnic boundary), then local cooperation will dominate. Similarly, this model is consistent with corruption as a lower-scale of cooperation undermining a higher scale^9,14,15^ and supports findings that lower-scale cooperation is more stable than a higher scale^25,31,32^. If an invasion by local cooperators is possible in the first place, then inevitably all other local groups will also convert to cooperating locally. This suggests that small-scale corruption erodes cooperation at higher scales and can lead to fracturing within a society. Because corruption as lower-scale cooperation within a large group will degrade the possibility of all group members working together, any society with some corruption risks descending into a wholly corrupt system where cooperation only occurs within local groups. Finally, the model is also related to research on parochial cooperation. Research shows that in-group favouritism leads to a preference for working with like-minded local group rather than a diverse global group^33–35^. These psychological mechanisms may be a proximate manifestation of the ultimate dynamics of overlapping scales of cooperation modelled here^36^.

Indirect reciprocity maintains cooperation at different cooperative scales. These multiple scales of indirect reciprocity protect cooperators from descending into full defection, but they also compete with and undermine one another. As such, under a range of conditions, societies are at risk of collapsing to lower scales of cooperation, which may help explain fracturing and corruption in previously cooperative societies as a result of resource constraints or slowed economic growth reducing payoffs at a larger societal scale.

## Methods

We consider a group of *n*_*g*_ individuals, which we will call our *global group*, subdivided into a groups of *n*_*l*_ individuals, which we will call *local groups*. Each member of the global group is assigned to one local group and each local group is independent from one another, which implies *n*_*g*_ = *a* · *n*_*l*_. Players begin by playing a Public Goods Game (PGG) with two investment pools, a global pool, which everyone has access to, and a local pool, which is unique to each local group and only that local group has access to. To cooperate in the PGG costs the cooperator *c*_*p*_ which becomes *b*_*g*_ if invested in the global pool or *b*_*l*_ if invested into the local pool. The relationship between *b*_*g*_ and *b*_*l*_ is undetermined, with it being possible for either game to be more beneficial. The PGG returns are then divided evenly amongst the respective groups. Players then play a Mutual Aid Game (MAG) within their local group. Here they are paired with one random member of their local group and with each partner they can choose to pay a cost c_*m*_ which yields their partner a benefit of *b*_*m*_. In turn, each player is also the receiving partner of another local group member, where they receive *b*_*m*_ when their partner pays the cost associated. Players know each other’s action from the previous round of the PGG and MAG, which will be used to determine how players act in future rounds. We also include an implementation error where players will accidentally do the opposite they intended in the MAG. The full list of parameters is listed in Table 1.

**Table 1:** Model parameters.

| Parameter | Meaning | Domain |
| --- | --- | --- |
| $a$ | Number of local groups | $> 0$ |
| $n_l$ | Number of people per local group | $> 0$ |
| $n_g$ | Number of people in the global group | $= a \cdot n_l$ |
| $c_p$ | Cost of contributing to either PGG | $> 0$ |
| $c_m$ | Cost of aiding in the MAG | $> 0$ |
| $b_l$ | Returns from local PGG contributions | $> c_p$ |
| $b_g$ | Returns from global PGG contributions | $> c_p$ |
| $b_m$ | Returns from being aided in the MAG | $> c_m$ |
| $e$ | Error rate | $0 < e < 1$ |

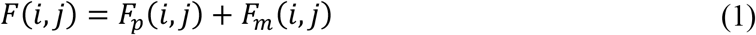

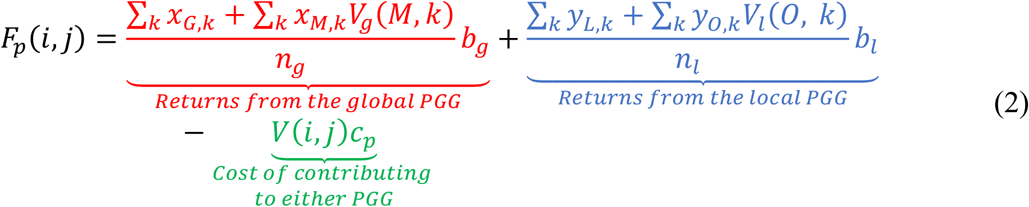

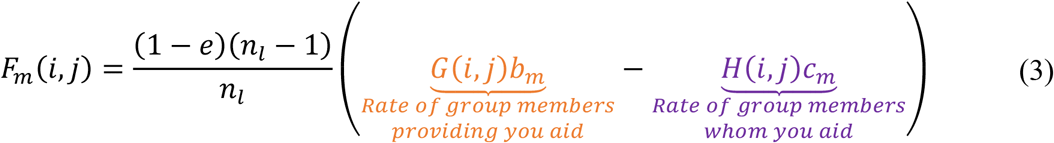

A player’s fitness (1) is broken down into two parts, PGG fitness (2) and MAG fitness (3). PGG fitness is comprised of returns from the global pool (in red), returns from the local pool (in blue), and the cost of contributing to either of the pools (in green). MAG fitness is comprised of aid received from other local group members (in orange) minus aid provided to other group members (in purple). This portion is determined by yours and others’ reputations. Players start with a positive reputation and lose their reputation if they fail to play in a manner others expect. That is, if I contribute to the global PGG but the others in my group expect me to contribute to the local PGG, then I fall out of good standing regardless of the fact I contributed to one of the PGGs.

The possible strategies for the PGG portion of the game are listed in Table 2 and for the MAG portion these are listed in Table 3. Note there are more strategies listed here than are discussed in the text’s body. The extra strategies are M and *O* in the PGG, which contribute to the global and local PGGs respectively but only if they’ve been aided in the MAG, and p in the MAG which aids all PGG cooperators regardless of which one that is. The results including *M, O*, and *p* are discussed in the appendix.

**Table 2.** Public Goods Game (PGG) strategies.

| PGG strategy | Description |
| --- | --- |
| $G$ | Always contributes to the global PGG pool |
| $L$ | Always contributes to the local PGG pool |
| $D$ | Does not contribute to either PGG |
| $M$ | Contributes to the global PGG pool, only if they received MAG aid |
| $O$ | Contributes to the local PGG pool, only if they received MAG aid |

**Table 3.**
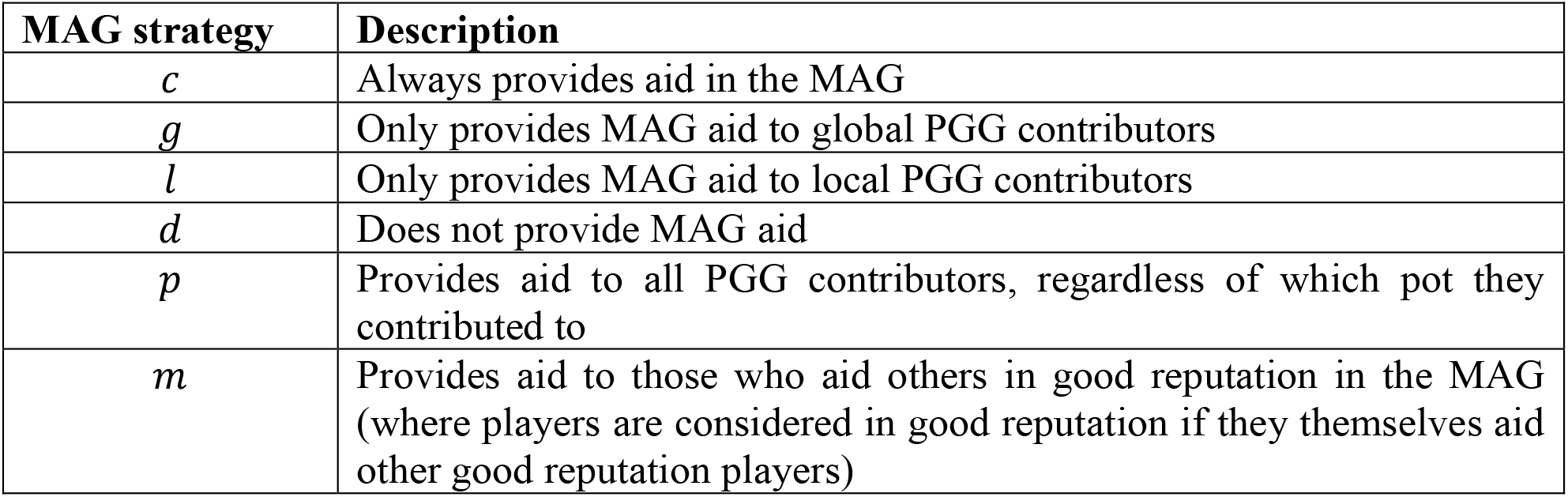
Mutual Aid Game (MAG) strategies.

Reputation is operationalized recursively. Players start with a good reputation and each round they can either lose their good reputation, or if it’s already lost then regain it. For a given round *n*, the percentage of players using strategy (*i, j*) with a good reputation, with some amount of variation due to a possible error *e* in implementing their strategy, is given by *W*_*n*_(*i, j*):

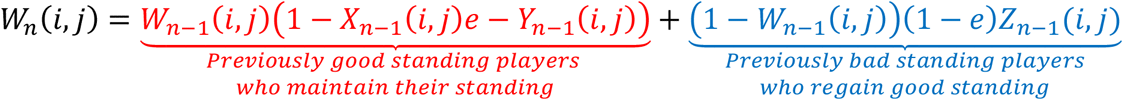

A player’s reputation in round *n* is dependent on their reputation in the previous round (*n* − 1) and their actions in the previous round (*n* − 1). In red we have the rate of players using strategy (*i, j*) who were in good standing previously and remain in good standing based on their actions in the MAG. In blue we have the rate of players using strategy (*i, j*) who were in bad standing previously but manage to regain a good standing based on their actions in the MAG. Here *X*(*i, j*) is the proportion of players in good MAG standing that players with strategy (*i, j*) give to, *Y*(*i, j*) is the proportion of players in good MAG standing that players with strategy (*i, j*) don’t give to, and *Z*(*i, j*) equals all players that players with strategy (*i, j*) give to, both in good and bad MAG standing. At equilibrium this equals:

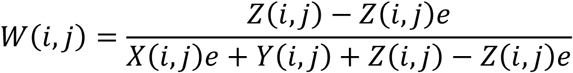

## Supporting information

Invasion Analysis Code

Invasion Analysis Results

## Data Availability

Methods for deriving results are described in greater detail within the Supplementary Information. Table of invasion results is also provided as Supplementary Information.

## Code Availability

The results of the invasion analysis were solved using Python SymPy symbolic mathematics computer algebra system (CAS) package. All code has been made available in the supplementary materials.

## Author Contributions

Eric Schnell and Michael Muthukrishna both conceived the model, discussed the model findings and wrote the manuscript. Eric Schnell developed and analyzed the model.

