## Supplementary Materials for "Indirect Reciprocity Undermines Indirect Reciprocity Destabilizing Large-Scale Cooperation"

October 24, 2023

### 1 Model Set up

We begin by writing the payoffs for each strategy. The strategies for the PGG are as follows:

| **Symbol** | **Behaviour** |
|---|---|
| $G$ | Contribute to Global PGG |
| $L$ | Contribute to Local PGG |
| $D$ | Defect from PGG |
| $M$ | Contribute to Global PGG only if you've been aided in the MAG |
| $O$ | Contribute to Local PGG only if you've been aided in the MAG |

Table 1: PGG Strategies

The strategies for the MAG are as follows:

| **Symbol** | **Behaviour** |
|---|---|
| $c$ | Always aid |
| $g$ | Aid if partner contributed to Global PGG |
| $l$ | Aid if partner contributed to Local PGG |
| $d$ | Always defect |
| $p$ | Aid those in good reputation based on PGG contributions |
| $m$ | Aid those in good reputation based on MAG contributions |
| $pm$ | Aid those in good reputation based on PGG contributions and MAG contributions |

Table 2: MAG Strategies

Thus, there are $5 \times 6 = 30$ different strategies. Note strategies $M$, $O$ and $p$ are omitted from the main text and only considered here. For our purposes we need to define 10 functions, as listed in table 3. To begin recall the payoff formulas as stated in the methods:

$$F(i,j) = F_p(i,j) + F_m(i,j) \tag{1}$$

$$F_p(i,j) = \frac{\sum_k x_{G,k} + \sum_k x_{M,k} V_g(M,k)}{n_g} b_g + \frac{\sum_k x_{L,k} + \sum_k x_{O,k} V_l(O,k)}{n_l} b_l - V(i,j) c_p \tag{2}$$

$$F_m(i,j) = \frac{(1-e)(n_l - 1)}{n_l} \left( G(i,j) b_m - H(i,j) c_m \right) \tag{3}$$

| **Function** | **Definition** |
|---|---|
| $F(i,j)$ | Fitness of player using strategy $(i,j)$ |
| $F_p(i,j)$ | Fitness derived from PGG of player using strategy $(i,j)$ |
| $F_m(i,j)$ | Fitness derived from MAG of player using strategy $(i,j)$ |
| $V_g(i,j)$ | Proportion of players using strategy $(i,j)$ who contribute to the global PGG |
| $V_l(i,j)$ | Proportion of players using strategy $(i,j)$ who contribute to the local PGG |
| $V(i,j)$ | Proportion of players using strategy $(i,j)$ who contribute to either PGG. Also defined as $V^g(i,j)+V^l(i,j)$ |
| $W(i,j)$ | Proportion of players using strategy $(i,j)$ who are in good standing in the MAG |
| $G(i,j)$ | Number of local group members who provide aid in the MAG to players using strategy $(i,j)$ |
| $H(i,j)$ | Number of local group members who players using strategy $(i,j)$ will provide aid to in the MAG |

Table 3: Model functions

Equations 2 and 3 both rely on defining more functions, specifically related to questions of reputation. Equation 2 relies on defining whether or not someone contributes to a public good (variations of $V$). Note equation 2 can be written in a more general form where all strategies are checked to see if they give contributions (instead of say only taking $V_g$ for strategies using $M$ in the PGG), but $V_g$ and $V_l$ will be either 1 or 0 for all strategies except $M$ and $O$, so other strategies can be excluded. Equation 3 relies on defining who a player receives aid from ($G(i,j)$) and who players provide aid to ($H(i,j)$), as determined by reputation.

Note when we run the analysis we assume $\frac{n_l-1}{n_l} \approx 1$ in equation 3. This makes the analysis easier to interpret without changing the insights.

We denote three different types of PGG standing, one in the global PGG (denoted by $V_g\left(i,j\right)$), another in the local PGG (denoted by $V_l\left(i,j\right)$) and finally an overall PGG standing (denoted by $V\left(i,j\right)$), where the overall PGG standing is the sum of these other two numbers ($V\left(i,j\right) = V_g\left(i,j\right) + V_l\left(i,j\right)$). PGG standing is defined for a given round $n$, denoted by a superscript, and is reliant upon MAG actions, specifically $W(i,j)$ which will be defined shortly. Note that the important variable of the function is the PGG strategy and we will explicitly define the result for each strategy.

$$
\begin{aligned}
V_g^n(G,j) &= 1; V_l^n(G,j) = 0 \\
V_g^n(L,j) &= 0; V_l^n(L,j) = 1 \\
V_g^n(D,j) &= 0; V_l^n(D,j) = 0 \\
V_g^n(M,j) &= \sum_K \Bigg( \frac{y_{K,c}W^{n-1}(K,c) + y_{K,g}W^{n-1}(K,g) + y_{K,p}W^{n-1}(K,p)}{n_l} \\
&\quad + \frac{y_{K,m}W^{n-1}(K,m)W^{n-1}(M,j) + y_{K,pm}W^{n-1}(K,pm)W^{n-1}(M,j)}{n_l} \Bigg) \\
V_l^n(M,j) &= 0 \\
V_g^n(O,j) &= 0 \\
V_l^n(O,j) &= \sum_K \Bigg( \frac{y_{K,c}W^{n-1}(K,c) + y_{K,g}W^{n-1}(K,g) + y_{K,p}W^{n-1}(K,p)}{n_l} \\
&\quad + \frac{y_{K,m}W^{n-1}(K,m)W^{n-1}(M,j) + y_{K,pm}W^{n-1}(K,pm)W^{n-1}(M,j)}{n_l} \Bigg)
\end{aligned} \tag{4}
$$

And now to define $W^n i,j$. We define this recursively where everyone begins in good standing, or $W^1(i,j) = 1$.

$$
W^n(i,j) = W^{n-1}(i,j)\left(1 - X(i,j)e - Y(i,j)\right) + \left(1 - W^{n-1}(i,j)\right)(1-e)\,Z(i,j) \tag{5}
$$

Here $X(i,j)$ is the proportion of players which contributed to the MAG and that players with strategy $(i,j)$ give to, $Y(i,j)$ is the proportion of players which contributed to the MAG that players with strategy $(i,j)$ don't give to, and $Z(i,j)$ equals all players that players with strategy $(i,j)$ give to, regardless of MAG . These will all be defined below, but first we can define $W(i,j)$ at equilibrium:

$$
\begin{aligned}
W(i,j) = W(i,j)\left(1 - X(i,j)e - Y(i,j)\right) + \left(1 - W(i,j)\right)(1-e)\,Z(i,j) \\
W(i,j) - W(i,j)\left(1 - X(i,j)e - Y(i,j)\right) + W(i,j)(1-e)\,Z(i,j) = (1-e)\,Z(i,j) \\
W(i,j)\left(1 - 1 + X(i,j)e + Y(i,j) + Z(i,j) - Z(i,j)e\right) = Z(i,j) - Z(i,j)e \\
W(i,j) = \frac{Z(i,j) - Z(i,j)e}{X(i,j)e + Y(i,j) + Z(i,j) - Z(i,j)e}
\end{aligned} \tag{6}
$$

$X(i,j)$, $Y(i,j)$ and $Z(i,j)$ are only dependent on MAG strategy and we will define their values for each of these.

For MAG strategy $c$:

$$
\begin{aligned}
X(i,c) &= \sum_{K,k} \frac{y_{K,k}W(K,k)}{n_l} \\
Y(i,c) &= 0 \\
Z(i,c) &= 1
\end{aligned}
$$

Given a small enough error rate, such that $e^2 \approx 0$, $W(i,c)$ can be further simplified to:

$$
W(i,c) = 1 - X(i,c)e
$$

For MAG strategy $g$:

$$X(i,g) = \sum_{K,k} \frac{y_{K,k} W(K,k) V_g(K,k)}{n_l}$$
$$Y(i,g) = \sum_{K,k} \frac{y_{K,k} W(K,k) \left(1 - V_g(K,k)\right)}{n_l}$$
$$Z(i,g) = \sum_{K,k} \frac{y_{K,k} V_g(K,k)}{n_l}$$

For MAG strategy $l$:

$$X(i,l) = \sum_{K,k} \frac{y_{K,k} W(K,k) V_l(K,k)}{n_l}$$
$$Y(i,l) = \sum_{K,k} \frac{y_{K,k} W(K,k) \left(1 - V_l(K,k)\right)}{n_l}$$
$$Z(i,l) = \sum_{K,k} \frac{y_{K,k} V_l(K,k)}{n_l}$$

For MAG strategy $d$:

$$X(i,d) = 0$$
$$Y(i,d) = \sum_{K,k} \frac{y_{K,k} W(K,k)}{n_l}$$
$$Z(i,d) = 0$$

Note that this means $W(i,d) = 0$.
For MAG strategy $p$:

$$X(i,p) = \sum_{K,k} \frac{y_{K,k} W(K,k) V(K,k)}{n_l}$$
$$Y(i,p) = \sum_{K,k} \frac{y_{K,k} W(K,k) \left(1 - V(K,k)\right)}{n_l}$$
$$Z(i,p) = \sum_{K,k} \frac{y_{K,k} V(K,k)}{n_l}$$

For MAG strategy $m$:

$$X(i,m) = \sum_{K,k} \frac{y_{K,k} W(K,k)}{n_l}$$
$$Y(i,m) = 0$$
$$Z(i,m) = \sum_{K,k} \frac{y_{K,k} W(K,k)}{n_l}$$

Note that $X(i,m) = Z(i,m)$, we can thus make the simplification $W(i,m) = 1 - e$.

For MAG strategy $pm$:

$$X(i,pm) = \sum_{K,k} \frac{y_{K,k} W(K,k) V(K,k)}{n_l}$$
$$Y(i,pm) = 0$$
$$Z(i,pm) = \sum_{K,k} \frac{y_{K,k} W(K,k) V(K,k)}{n_l}$$

Note that $X(i,pm) = Z(i,pm)$, we can thus make the simplification $W(i,pm) = 1 - e$.

Note that these functions are circular and dependent on themselves, with $W(i,j)$ being dependent on $X(i,j)$, $Y(i,j)$ and $Z(i,j)$ and these being dependent on $W(i,j)$. To break this cycle, we assume all $W(i,j)$ in one of these functions is 1, except for MAG defectors in which case it is 0.

We conduct the invasion analysis by comparing resident fitness to invader fitness. This was conduct in Python SymPy symbolic mathematics computer algebra system (CAS) package and the code for running the analysis can be found in the supplementary materials. The full results of the analysis are compiled in an excel spreadsheet, also found in the supplementary materials. Note some of the results are re-evaluated by hand because SymPy was unable to tell whether it was greater or less than 0. For example, $(G,l)$ is able to invade $(G,c)$ in a single invader model when $cm \cdot (1-e) > 0$. Based on our assumptions about our parameters ($cm > 0$ and $0 < e < 1$) we know this invasion will always occur, but SymPy was unable to detect it as such so we make this change in the results.
