## Supplementary material for "Indirect Reciprocity Undermines Indirect Reciprocity Destabilizing Large-Scale Cooperation": Invasion Analysis Code: Local_global_analysis_v01.py

```
from sympy import *
import time
from datetime import timedelta
import pandas as pd

# Establish resident and invader populations
# First do so for the local group being invaded
resident_prop_a = symbols('resident_prop_a')
invader_prop_a = 1 - resident_prop_a
# Then for all other groups
resident_prop_other = symbols('resident_prop_other')
invader_prop_other = 1 - resident_prop_other

# We use relatedness to determine how many residents and invaders there are. This could be included in the proportions
# instead but here we only set proportions to 1 or 0 and modify it using relatedness
relatedness = symbols('relatedness')
resident_expect_a = (1 - relatedness) * resident_prop_a
invader_expect_a = relatedness + (1 - relatedness) * invader_prop_a
resident_expect_other = resident_prop_other
invader_expect_other = invader_prop_other

# Create resident and invader playstyles as global variables
resident_playstyle = (None, None)
invader_playstyle = (None, None)

# Costs and benefits to games
cp = symbols('cp', positive=True)
cm = symbols('cm', positive=True)

# Enforce benefits to be greater than costs
bg_minus_cp = symbols('bg_minus_cp', positive=True)
bl_minus_cp = symbols('bl_minus_cp', positive=True)
bm_minus_cm = symbols('bm_minus_cm', positive=True)

# Make two forms of all parameters, a complicated one and a simple one
# Complicated form carries more information but sympy has an easier time telling which is greater
bg_comp = bg_minus_cp + cp
bl_comp = bl_minus_cp + cp
bm_comp = bm_minus_cm + cm

# Simple form is more legible
bg_simp = symbols('bg_simp', positive=True)
bl_simp = symbols('bl_simp', positive=True)
bm_simp = symbols('bm_simp', positive=True)

# Number of players
d_one = symbols('d_one', positive=True, integer=True)
num_groups = symbols('num_groups', positive=True, integer=True)

# This ensures that both nl is greater than 1 and that ng is greater than nl
nl_comp = 1 + d_one
ng_comp = nl_comp * num_groups

nl_simp = symbols('nl_simp', positive=True, integer=True)
ng_simp = nl_simp * num_groups

# Error rate
# This ensures that e is between 0 and 1
t = symbols('t', nonnegative=True)
e_comp = t / (1 + t)

# This is a simpler and more legible form of e
e_simp = symbols('e_simp', positive=True)

# List all playstyles
PGG_playstyles = ['G', 'L', 'D', 'M', 'O']
MAG_playstyles = ['c', 'g', 'l', 'd', 'p', 'm', 'pm']

# Store values
vals = {}


def give_to(playstyle):
    """
    Returns a list of playstyles that a player will give to in the MAG, given their own 'playstyle'.

    :param playstyle: Player's playstyle
    :type playstyle: (string, string)
    :return: list of playsyles
    :rtype: list
    """
    give_list = []
    if playstyle[1] == 'c':
        give_list += [(a, b) for a in PGG_playstyles for b in MAG_playstyles]
        # sm doesn't give to every combo but this will be checked later
    elif playstyle[1] == 'g':
        give_list += [('G', b) for b in MAG_playstyles]
        give_list += [('M', b) for b in MAG_playstyles]
    elif playstyle[1] == 'l':
        give_list += [('L', b) for b in MAG_playstyles]
        give_list += [('O', b) for b in MAG_playstyles]
    elif playstyle[1] == 'p':
        give_list += [(a, b) for a in ['G', 'L', 'M', 'O'] for b in MAG_playstyles]
    elif playstyle[1] == 'm':
        give_list += [(a, 'c') for a in PGG_playstyles]
        give_list += [(a, 'm') for a in PGG_playstyles]
        give_list += [(a, 'p') for a in ['G', 'L', 'M', 'O']]
        give_list += [('G', 'g'), ('M', 'g'), ('L', 'l'), ('O', 'l')]
        give_list += [(a, 'pm') for a in ['G', 'L', 'M', 'O']]
    elif playstyle[1] == 'pm':
        give_list += [(a, 'c') for a in ['G', 'L', 'M', 'O']]
        give_list += [(a, 'm') for a in ['G', 'L', 'M', 'O']]
        give_list += [('G', 'g'), ('M', 'g'), ('L', 'l'), ('O', 'l')]
        give_list += [(a, 'p') for a in ['G', 'L', 'M', 'O']]
        give_list += [(a, 'pm') for a in ['G', 'L', 'M', 'O']]
    return give_list


def give_to_plus_defect(playstyle):
    """
    Returns a list of playstyles that a player will give to in the MAG, given their own 'playstyle'. This function is
    used to determine roughly a player's standing. As a result include defectors as failing to give to these does not
    reduce standing.

    :param playstyle: Player's playstyle
    :type playstyle: (string, string)
    :return: list of playsyles
    :rtype: list
    """
    give_list = give_to(playstyle)
    # This may include duplicates, but functionally does not change outcome.
    give_list += [(a, 'd') for a in PGG_playstyles]
    return give_list


def receive_from_stand(playstyle, within_a=True):
    """
    Returns what proportion of local players will provide aid to someone using strategy 'playstyle'.

    :param playstyle: Player's playstyle
    :type playstyle: (string, string)
    :param within_a: 'True' to calculate within invaded group and 'False' to calculate in full resident group
    :type within_a: bool, optional
    :return: Proportion of players that playstyle receives from
    :rtype: float
    """
    if within_a:
        resident_expect = resident_expect_a
        invader_expect = invader_expect_a
    else:
        resident_expect = resident_expect_other
        invader_expect = invader_expect_other
    x = 0

    # Amount of aid received from residents
    if playstyle in give_to(resident_playstyle):
        weight = MAG_standing_binary(resident_playstyle, within_a)
        x += weight * resident_expect

    # Amount of aid received from invaders
    if playstyle in give_to(invader_playstyle):
        weight = MAG_standing_binary(invader_playstyle, within_a)
        x += weight * invader_expect

    return x


def MAG_standing_approx(playstyle, simple=True, within_a=True):
    """
    Returns the approximate MAG standing at equilibrium for players with the strategy 'playstyle'. Details for deriving
    this equation appear in the appendix. Results are stored in a dictionary and can be retrieved for faster completion.

    Note: Based on P-B 2004, fall out of standing for failing to help player in good standing during MAG, but regain
    standing by helping any player during MAG, even those in bad standing.

    :param playstyle: Player's playstyle
    :type playstyle: (string, string)
    :param simple: 'True' to use simple definitions and 'False' to use complex definitions
    :type simple: bool, optional
    :param within_a: 'True' to calculate within invaded group and 'False' to calculate in full resident group
    :type within_a: bool, optional
    :return: MAG standing of the player
    :rtype: float
    """
    # Sets which definition of e will be used
    if simple:
        e = e_simp
    else:
        e = e_comp

    # Standing is dependent on players in good standing who you aid, players in good standing who you don't aid and all
    # players you aid.
    give_stand = give_to_standing(playstyle, simple, within_a)
    give_all = give_to_all(playstyle, simple, within_a)
    not_give_stand = not_give_to_standing(playstyle, simple, within_a)

    top = give_all - give_all * e
    bottom = give_stand * e + not_give_stand + give_all - give_all * e

    if top.subs(vals) == 0 and bottom.subs(vals) == 0:
        # Should only happen when all players are defectors because then they give to no one but there's no one in good
        # standing who should've been given to anyway
        stand = 0
    else:
        stand = (top / bottom).subs(vals)

    return stand


def give_to_standing(playstyle, simple=True, within_a=True):
    """
    Returns the proportion of local players in good standing that players using the strategy 'playstyle' will aid in
    the MAG.

    Because of circular nature of logic, uses a simplified logic assumption for determining other standings, as
    detailed in MAG_standing_binary.

    :param playstyle: Player's playstyle
    :type playstyle: (string, string)
    :param simple: 'True' to use simple definitions and 'False' to use complex definitions
    :type simple: bool, optional
    :param within_a: 'True' to calculate within invaded group and 'False' to calculate in full resident group
    :type within_a: bool, optional
    :return: standing of players being given to
    :rtype: float
    """
    if within_a:
        resident_expect = resident_expect_a
        invader_expect = invader_expect_a
    else:
        resident_expect = resident_expect_other
        invader_expect = invader_expect_other

    result = 0
    # 'c' and 'm' aid all good standing players
    if playstyle[1] == 'c' or playstyle[1] == 'm':
        result += resident_expect * MAG_standing_binary(resident_playstyle, within_a)
        result += invader_expect * MAG_standing_binary(invader_playstyle, within_a)
    # 'g' aid all good standing players who contributed to global PGG
    elif playstyle[1] == 'g':
        result += (resident_expect * PGG_standing_approx(resident_playstyle, simple, within_a)[0] *
                   MAG_standing_binary(resident_playstyle, within_a))
        result += (invader_expect * PGG_standing_approx(invader_playstyle, simple, within_a)[0] *
                   MAG_standing_binary(invader_playstyle, within_a))
    # 'l' aid all good standing players who contributed to local PGG
    elif playstyle[1] == 'l':
        result += (resident_expect * PGG_standing_approx(resident_playstyle, simple, within_a)[1] *
                   MAG_standing_binary(resident_playstyle, within_a))
        result += (invader_expect * PGG_standing_approx(invader_playstyle, simple, within_a)[1] *
                   MAG_standing_binary(invader_playstyle, within_a))
    # 'p' and 'pm' aid all good standing players who contributed to global PGG
    elif playstyle[1] == 'p' or playstyle[1] == 'pm':
        result += (resident_expect * PGG_standing_approx(resident_playstyle, simple, within_a)[2] *
                   MAG_standing_binary(resident_playstyle, within_a))
        result += (invader_expect * PGG_standing_approx(invader_playstyle, simple, within_a)[2] *
                   MAG_standing_binary(invader_playstyle, within_a))

    return result


def not_give_to_standing(playstyle, simple=True, within_a=True):
    """
    Returns the proportion of local players in good MAG standing that players using the strategy 'playstyle' will not
    aid in the MAG.

    Because of circular nature of logic, uses a simplified logic assumption for determining other standings, as
    detailed in MAG_standing_binary.

    :param playstyle: Player's playstyle
    :type playstyle: (string, string)
    :param simple: 'True' to use simple definitions and 'False' to use complex definitions
    :type simple: bool, optional
    :param within_a: 'True' to calculate within invaded group and 'False' to calculate in full resident group
    :type within_a: bool, optional
    :return: standing of players not being given to
    :rtype: float
    """
    if within_a:
        resident_expect = resident_expect_a
        invader_expect = invader_expect_a
    else:
        resident_expect = resident_expect_other
        invader_expect = invader_expect_other

    result = 0
    if playstyle[1] == 'd':
        result += resident_expect * MAG_standing_binary(resident_playstyle, within_a)
        result += invader_expect * MAG_standing_binary(invader_playstyle, within_a)
    elif playstyle[1] == 'g':
        result += (resident_expect * (1 - PGG_standing_approx(resident_playstyle, simple, within_a)[0])
                   * MAG_standing_binary(resident_playstyle, within_a))
        result += (invader_expect * (1 - PGG_standing_approx(invader_playstyle, simple, within_a)[0])
                   * MAG_standing_binary(invader_playstyle, within_a))
    elif playstyle[1] == 'l':
        result += (resident_expect * (1 - PGG_standing_approx(resident_playstyle, simple, within_a)[1])
                   * MAG_standing_binary(resident_playstyle, within_a))
        result += (invader_expect * (1 - PGG_standing_approx(invader_playstyle, simple, within_a)[1])
                   * MAG_standing_binary(invader_playstyle, within_a))
    elif playstyle[1] == 'p':
        result += (resident_expect * (1 - PGG_standing_approx(resident_playstyle, simple, within_a)[2])
                   * MAG_standing_binary(resident_playstyle, within_a))
        result += (invader_expect * (1 - PGG_standing_approx(invader_playstyle, simple, within_a)[2])
                   * MAG_standing_binary(invader_playstyle, within_a))

    return result


def give_to_all(playstyle, simple=True, within_a=True):
    """
    Returns the proportion of local players that players using the strategy 'playstyle' will aid in the MAG, regardless
    of their standing.

    Because of circular nature of logic, uses a simplified logic assumption for determining other standings, as
    detailed in MAG_standing_binary.

    :param playstyle: Player's playstyle
    :type playstyle: (string, string)
    :param simple: 'True' to use simple definitions and 'False' to use complex definitions
    :type simple: bool, optional
    :param within_a: 'True' to calculate within invaded group and 'False' to calculate in full resident group
    :type within_a: bool, optional
    :return: standing of players not being given to
    :rtype: float
    """
    if within_a:
        resident_expect = resident_expect_a
        invader_expect = invader_expect_a
    else:
        resident_expect = resident_expect_other
        invader_expect = invader_expect_other

    result = 0
    if playstyle[1] == 'c':
        result = 1
    elif playstyle[1] == 'g':
        result += resident_expect * PGG_standing_approx(resident_playstyle, simple, within_a)[0]
        result += invader_expect * PGG_standing_approx(invader_playstyle, simple, within_a)[0]
    elif playstyle[1] == 'l':
        result += resident_expect * PGG_standing_approx(resident_playstyle, simple, within_a)[1]
        result += invader_expect * PGG_standing_approx(invader_playstyle, simple, within_a)[1]
    elif playstyle[1] == 'p':
        result += resident_expect * PGG_standing_approx(resident_playstyle, simple, within_a)[2]
        result += invader_expect * PGG_standing_approx(invader_playstyle, simple, within_a)[2]
    elif playstyle[1] == 'm':
        result += resident_expect * MAG_standing_binary(resident_playstyle, within_a)
        result += invader_expect * MAG_standing_binary(invader_playstyle, within_a)
    elif playstyle[1] == 'pm':
        result += (resident_expect * PGG_standing_approx(resident_playstyle, simple, within_a)[2]
                   * MAG_standing_binary(resident_playstyle, within_a))
        result += (invader_expect * PGG_standing_approx(invader_playstyle, simple, within_a)[2]
                   * MAG_standing_binary(invader_playstyle, within_a))

    return result


def MAG_standing_binary(playstyle, within_a=True):
    """
    Returns a binary approximation for the standing of a player using the strategy 'playstyle'. This approximation is
    used to avoid infinite recursion while calculating MAG standing. Calculated by saying players are in good standing
    for giving to anyone or either giving or not giving to defectors. Except for defectors who always have bad standing.

    :param playstyle: Player's playstyle
    :type playstyle: (string, string)
    :param within_a: 'True' to calculate within invaded group and 'False' to calculate in full resident group
    :type within_a: bool, optional
    :return: binary standing
    :rtype: float
    """
    if within_a:
        resident_expect = resident_expect_a
        invader_expect = invader_expect_a
    else:
        resident_expect = resident_expect_other
        invader_expect = invader_expect_other

    result = 0

    if playstyle[1] == 'd':
        return result

    if resident_playstyle in give_to_plus_defect(playstyle):
        result += resident_expect
    if invader_playstyle in give_to_plus_defect(playstyle):
        result += invader_expect

    return result


def PGG_standing_approx(playstyle, simple=True, within_a=True):
    """
    Returns the approximate PGG standing at equilibrium for players with the strategy 'playstyle'.

    :param playstyle: Player's playstyle
    :type playstyle: (string, string)
    :param simple: 'True' to use simple definitions and 'False' to use complex definitions
    :type simple: bool, optional
    :param within_a: 'True' to calculate within invaded group and 'False' to calculate in full resident group
    :type within_a: bool, optional
    :return: tuple of global PGG standing, local PGG standing, and largest PGG standing
    :rtype: tuple
    """
    if simple:
        e = e_simp
    else:
        e = e_comp

    pair = (None, None)
    if playstyle[0] == 'D':
        pair = (0, 0)  # 'D' never gives
    elif playstyle[0] == 'G':
        pair = (1, 0)  # 'G' always gives to the global PGG
    elif playstyle[0] == 'L':
        pair = (0, 1)  # 'L' always gives to the local PGG
    elif playstyle[0] == 'M':
        # Assume that players have an 'e' chance to fail to provide aid when they should have, so even if all the
        # players want to provide aid to players using 'playstyle' only 1 - e actually will, and those who didn't
        # receive any aid will not participate in the PGG.
        stand = receive_from_stand(playstyle, within_a) * (1 - e)
        pair = (stand, 0)  # 'M' only gives to the global PGG if they received MAG aid
    elif playstyle[0] == 'O':
        stand = receive_from_stand(playstyle, within_a) * (1 - e)
        pair = (0, stand)  # 'O' only gives to the local PGG if they received MAG aid

    # Pull out the higher PGG standing for future ease
    # Can't use max because these results are sympy equations
    if pair[0] == 0:
        answer = pair[1]
    else:
        answer = pair[0]

    return pair[0], pair[1], answer


def fitness_approx(playstyle, simple=True):
    """
    Returns the approximate fitness of players with strategy 'playstyle'. Players receive PGG returns according to the
     number of players who give to the global and local PGG, minus how much they give to the PGG. Players receive MAG
     returns according to the number of players who will aid them minus how much they aid in the MAG.

    :param playstyle: Player's playstyle
    :type playstyle: (string, string)
    :param simple: 'True' to use simple definitions and 'False' to use complex definitions
    :type simple: bool, optional
    :return: Player fitness
    :rtype: float
    """
    # Sets which definition of e, ng, and nl will be used
    if simple:
        e = e_simp
        ng = ng_simp
        nl = nl_simp
        bg = bg_simp
        bl = bl_simp
        bm = bm_simp
    else:
        e = e_comp
        ng = ng_comp
        nl = nl_comp
        bg = bg_comp
        bl = bl_comp
        bm = bm_comp

    pay_global = 0
    pay_local = 0

    pay_global += resident_expect_a * nl * PGG_standing_approx(resident_playstyle, simple, True)[0]
    pay_global += invader_expect_a * nl * PGG_standing_approx(invader_playstyle, simple, True)[0]
    pay_global += (resident_expect_other * (num_groups - 1) * nl
                   * PGG_standing_approx(resident_playstyle, simple, False)[0])
    pay_global += (invader_expect_other * (num_groups - 1) * nl
                   * PGG_standing_approx(invader_playstyle, simple, False)[0])
    pay_local += resident_expect_a * nl * PGG_standing_approx(resident_playstyle, simple, True)[1]
    pay_local += invader_expect_a * nl * PGG_standing_approx(invader_playstyle, simple, True)[1]

    reward_global = bg * pay_global / ng
    reward_local = bl * pay_local / nl

    cost_PGG = cp * PGG_standing_approx(playstyle, simple, True)[2]

    reward_PGG = reward_global + reward_local - cost_PGG

    benefit_MAG = bm * total_amount_received(playstyle, simple, True)

    cost_MAG = cm * cost_paid(playstyle, simple, True)

    # For simplification, we omit '* (nl - 1) * (1 / nl)' as this doesn't change the result and makes it easier to
    # interpret
    reward_MAG = (benefit_MAG - cost_MAG) * (1 - e)

    fit = reward_PGG + reward_MAG

    return fit.subs(vals)


def total_amount_received(playstyle, simple=True, within_a=True):
    """
    Returns the amount a player with strategy 'playstyle' receives from the MAG.

    :param playstyle: Player's playstyle
    :type playstyle: (string, string)
    :param simple: 'True' to use simple definitions and 'False' to use complex definitions
    :type simple: bool, optional
    :param within_a: 'True' to calculate within invaded group and 'False' to calculate in full resident group
    :type within_a: bool, optional
    :return: Amount received
    :rtype: float
    """
    if within_a:
        resident_expect = resident_expect_a
        invader_expect = invader_expect_a
    else:
        resident_expect = resident_expect_other
        invader_expect = invader_expect_other

    result = 0
    res_weight = amount_received(playstyle, resident_playstyle, simple, within_a)
    inv_weight = amount_received(playstyle, invader_playstyle, simple, within_a)

    result += resident_expect * res_weight
    result += invader_expect * inv_weight

    return result


def amount_received(receiver_playstyle, giver_playstyle, simple=True, within_a=True):
    """
    Returns the total amount a player with strategy 'receiver_playstyle' receives in the MAG from a player with
    strategy 'giver_playstyle'.

    :param receiver_playstyle: Receiving player's playstyle
    :type receiver_playstyle: (string, string)
    :param giver_playstyle: Giving player's playstyle
    :type giver_playstyle: (string, string)
    :param simple: 'True' to use simple definitions and 'False' to use complex definitions
    :type simple: bool, optional
    :param within_a: 'True' to calculate within invaded group and 'False' to calculate in full resident group
    :type within_a: bool, optional
    :return: Amount received
    :rtype: float
    """
    amount = 0

    if giver_playstyle[1] == 'c':
        amount = 1
    elif giver_playstyle[1] == 'g':
        amount = PGG_standing_approx(receiver_playstyle, simple, within_a)[0]
    elif giver_playstyle[1] == 'l':
        amount = PGG_standing_approx(receiver_playstyle, simple, within_a)[1]
    elif giver_playstyle[1] == 'p':
        amount = PGG_standing_approx(receiver_playstyle, simple, within_a)[2]
    elif giver_playstyle[1] == 'm':
        amount = MAG_standing_approx(receiver_playstyle, simple, within_a)
    elif giver_playstyle[1] == 'pm':
        amount = (PGG_standing_approx(receiver_playstyle, simple, within_a)[2]
                  * MAG_standing_approx(receiver_playstyle, simple, within_a))

    return amount


def cost_paid(playstyle, simple=True, within_a=True):
    """
    Returns the amount a player with strategy 'playstyle' would give in the MAG.

    :param playstyle: Player's playstyle
    :type playstyle: (string, string)
    :param simple: 'True' to use simple definitions and 'False' to use complex definitions
    :type simple: bool, optional
    :param within_a: 'True' to calculate within invaded group and 'False' to calculate in full resident group
    :type within_a: bool, optional
    :return: Cost paid
    :rtype: float
    """
    if within_a:
        resident_expect = resident_expect_a
        invader_expect = invader_expect_a
    else:
        resident_expect = resident_expect_other
        invader_expect = invader_expect_other

    result = 0
    res_weight = 0
    inv_weight = 0

    if playstyle[1] == 'c':
        res_weight = 1
        inv_weight = 1
    elif playstyle[1] == 'g':
        res_weight = PGG_standing_approx(resident_playstyle, simple, within_a)[0]
        inv_weight = PGG_standing_approx(invader_playstyle, simple, within_a)[0]
    elif playstyle[1] == 'l':
        res_weight = PGG_standing_approx(resident_playstyle, simple, within_a)[1]
        inv_weight = PGG_standing_approx(invader_playstyle, simple, within_a)[1]
    elif playstyle[1] == 'p':
        res_weight = PGG_standing_approx(resident_playstyle, simple, within_a)[2]
        inv_weight = PGG_standing_approx(invader_playstyle, simple, within_a)[2]
    elif playstyle[1] == 'm':
        res_weight = MAG_standing_approx(resident_playstyle, simple, within_a)
        inv_weight = MAG_standing_approx(invader_playstyle, simple, within_a)
    elif playstyle[1] == 'pm':
        res_weight = PGG_standing_approx(resident_playstyle, simple, within_a)[2] * MAG_standing_approx(
            resident_playstyle, simple, within_a)
        inv_weight = PGG_standing_approx(invader_playstyle, simple, within_a)[2] * MAG_standing_approx(
            invader_playstyle, simple, within_a)

    result += resident_expect * res_weight
    result += invader_expect * inv_weight

    return result


if __name__ == '__main__':
    # Create all total (PGG, MAG) strategy combinations
    combos = []
    for a in PGG_playstyles:
        for b in MAG_playstyles:
            combos.append("{}".format((a, b)))

    # Create dataframes for tracking invasion criteria and fitnesses
    df_inv = pd.DataFrame(columns=combos, index=combos)
    df_fit = pd.DataFrame(columns=combos, index=combos)

    # Script used to calculate all possible resident-invader combinations
    runtime = time.time()

    # Assumptions
    # vals[resident_prop_a] = 1 for single or multiple invaders
    # vals[resident_prop_a] = 0 for group invaders
    # vals[relatedness] = 0 for single invader
    # vals[relatedness] = 1 for group invaders
    # Comment out vals[relatedness] line for multiple invaders

    vals[resident_prop_a] = 1
    vals[resident_prop_other] = 1
    vals[relatedness] = 0.5

    # Uncomment these lines to have no error
    # vals[e_simp] = 0
    # vals[t] = 0

    # Uncomment these lines to run for a specific set of parameters
    # vals[cp] = 1
    # vals[cm] = 1
    # vals[bm_simp] = 3
    # vals[num_groups] = 5
    # vals[bg_simp] = 3
    # vals[bl_simp] = 2
    # vals[e_simp] = 0.05
    # vals[nl_simp] = 5

    # Go through all combinations of residents and invaders
    for pgg_res in PGG_playstyles:
        for mag_res in MAG_playstyles:
            resident_playstyle = (pgg_res, mag_res)
            invader_playstyle = (pgg_res, mag_res)

            print("resident is {}".format(resident_playstyle))

            resident_comp = fitness_approx(resident_playstyle, False)
            resident_simp = fitness_approx(resident_playstyle, True)
            row_1 = {}
            row_2 = {}
            print('{} resident fitness = {}\n'.format(resident_playstyle, resident_simp))

            for pgg_inv in PGG_playstyles:
                for mag_inv in MAG_playstyles:
                    invader_playstyle = (pgg_inv, mag_inv)

                    mutant_comp = fitness_approx(invader_playstyle, False)
                    mutant_simp = fitness_approx(invader_playstyle, True)

                    # If mutant and resident fitness are equal then no invasion
                    if simplify(mutant_comp - resident_comp == 0):
                        invasion = None
                    else:
                        invasion = simplify(mutant_simp - resident_simp > 0)

                    row_1["{}".format(invader_playstyle)] = invasion
                    row_2["{}".format(invader_playstyle)] = mutant_simp
                    print('{} invasion fitness = {}'.format(invader_playstyle, mutant_simp))
                    print('{} invasion: {}\n'.format(invader_playstyle, invasion))

            df_inv.loc['{}'.format(resident_playstyle)] = row_1
            df_fit.loc['{}'.format(resident_playstyle)] = row_2

    elapsed = time.time() - runtime
    time_minutes = timedelta(seconds=elapsed)
    print("\n{}\n".format(time_minutes))
    runtime = time.time()

    df_inv.replace({False: -1, None: 0, True: 1}, inplace=True)

    # Save results
    # df_inv.to_csv('')
    # df_fit.to_csv('')
```
